# Modeling Flow in an *In Vitro* Anatomical Cerebrovascular Model with Experimental Validation

**DOI:** 10.1101/2023.01.13.523948

**Authors:** Saurabh Bhardwaj, Brent A. Craven, Jacob E. Sever, Francesco Costanzo, Scott D. Simon, Keefe B. Manning

**Affiliations:** Department of Biomedical Engineering, Pennsylvania State University, University Park, PA, USA; Office of Science and Engineering Laboratories, Center for Devices and Radiological Health, U.S. Food and Drug Administration, Silver Spring, MD, USA; Department of Engineering Science and Mechanics, Pennsylvania State University, University Park, PA, USA; Department of Neurosurgery, Penn State Hershey Medical Center, Hershey, PA, USA; Department of Surgery, Penn State Hershey Medical Center, Hershey, PA, USA

**Author notes:** **Corresponding Authors:** BA Craven, KB Manning.

**Keywords:** Cerebrovascular model, Cerebral blood flow, Image based modeling, Acute ischemic stroke

## Abstract

Acute ischemic stroke (AIS) is a leading cause of mortality that occurs when an embolus becomes lodged in the cerebral vasculature and obstructs blood flow in the brain. The severity of AIS is determined by the location and how extensively emboli become lodged, which are dictated in large part by the cerebral flow and the dynamics of embolus migration which are difficult to measure *in vivo* in AIS patients. Computational fluid dynamics (CFD) can be used to predict the patient-specific hemodynamics and embolus migration and lodging in the cerebral vasculature to better understand the underlying mechanics of AIS. To be relied upon, however, the computational simulations must be verified and validated. In this study, a realistic *in vitro* experimental model and a corresponding computational model of the cerebral vasculature are established that can be used to investigate flow and embolus migration and lodging in the brain. First, the *in vitro* anatomical model is described, including how the flow distribution in the model is tuned to match physiological measurements from the literature. Measurements of pressure and flow rate for both normal and stroke conditions were acquired and corresponding CFD simulations were performed and compared with the experiments to validate the flow predictions. Overall, the CFD simulations were in relatively close agreement with the experiments, to within ±7% of the mean experimental data with many of the CFD predictions within the uncertainty of the experimental measurement. This work provides an *in vitro* benchmark data set for flow in a realistic cerebrovascular model and is a first step towards validating a computational model of AIS.

## 1. Introduction

Stroke is one of the leading causes of mortality worldwide (over 6.5 million deaths/year) [1], with acute ischemic stroke (AIS) accounting for 87% of the total stroke mortalities [2]. AIS is a life-threatening medical condition that occurs when an embolus becomes lodged in the cerebral vasculature and obstructs blood flow in the brain. The available literature contains substantial research that has been performed to investigate cerebral blood flow (e.g., see [3-9]). A main focus of prior research has been to study regional cerebral blood flow, which forms the basis for identifying the causes of several diseases. Generally, the severity of AIS is determined by the location and how extensively emboli become lodged in the cerebral vasculature. It is extremely challenging, however, to measure the flow rate, pressure, and embolus migration in AIS patients. This level of information, though, is needed to explore the underlying causes and treatment of AIS.

The size and location of blood emboli in the cerebral vasculature have been the subject of several studies [10-16]. The middle cerebral artery (MCA) is the location where emboli most frequently lodge [13]. Measurements of the distribution of cerebral emboli within various arterial branches have been reported. A recent investigation using rats [14] showed that emboli shape and composition have a significant role in determining the severity of brain damage after they reach the cerebral vasculature. Using idealized Y-bifurcation geometries, Pollanen [15] and, more recently, Bushi et al. [16] conducted experiments on embolic particle migration. Beyond what volumetric flow patterns may imply, their research revealed a bias in the distribution of larger particles into the larger branching arteries like the common carotid artery (CCA), internal carotid artery (ICA), and the MCA. In their studies on embolus transport in the cerebral arteries, Chung et al. [10] used a more accurate anatomical model of the Circle of Willis and proposed a similar tendency of larger particles migrating toward the larger branching vessels. The distribution fraction of emboli across major arterial branches has been quantified in these experimental investigations corresponding to the associated volumetric flow patterns. But, they generally neglect the smaller branches along with the aortic arch in the anatomical model that can significantly influence the flow and embolus transport.

Computational fluid dynamics (CFD) can be used to predict patient-specific hemodynamics and embolus migration and lodging in the cerebral vasculature to better understand the underlying mechanics of AIS and to improve recanalization methods. Patient-specific CFD has become widely used for simulating blood flow in the cardiovascular system [17]; examples include evaluating the hemodynamics of healthy and diseased blood vessels [18-19], assisting in the design and assessment of vascular medical devices [20-21], planning vascular surgeries, and predicting the results of interventions [22-23]. To be relied upon, however, the computational simulations must be verified and validated, which is difficult to achieve *in vivo*, particularly for embolus migration. Realistic *in vitro* models can be used to provide a rich data set for CFD validation. For example, researchers at the U.S. Food and Drug Administration (FDA) and collaborators have developed benchmark models of a simplified nozzle, a centrifugal blood pump, and a vascular model of the inferior vena cava and have acquired flow measurements that are widely used for CFD validation [24-29].

The objective of this study is to establish an *in vitro* experimental model and a corresponding computational model of the cerebral vasculature that can be used to investigate flow and embolus migration and lodging in the brain. Levaraging a representative anatomical cerebrovascular model, computational simulations are performed with particular focus on comparing with experimental pressure and flow rate measurements. In addition to normal physiological flow conditions, the effect of arterial embolus occlusion in the MCA is considered to emulate a stroke to investigate its influence on cerebral blood flow.

## 2. Methods

### 2.1. Cerebrovascular model

A representative anatomical model of the aorta and cerebral vasculature was used in this study (Fig. 1). The model was designed and fabricated by United Biologics (Irvine, CA, USA) based on patient medical image data from several sources (e.g., the NIH Visible Human Project, patient-specific CT data). The *in vitro* model (Fig. 1) is made of silicone with an elastic modulus of 3.1-3.4 N/mm^2^, which is representative of human arteries. The entire model includes the aorta, common carotid arteries, internal and external carotid arteries, axillary arteries, middle cerebral arteries, and anterior cerebral arteries. In addition, a corresponding computational model of the *in vitro* model was reconstructed from high-resolution 3D micro computed tomography (μ-CT) scans (Fig. 2a).

**Figure 1.**
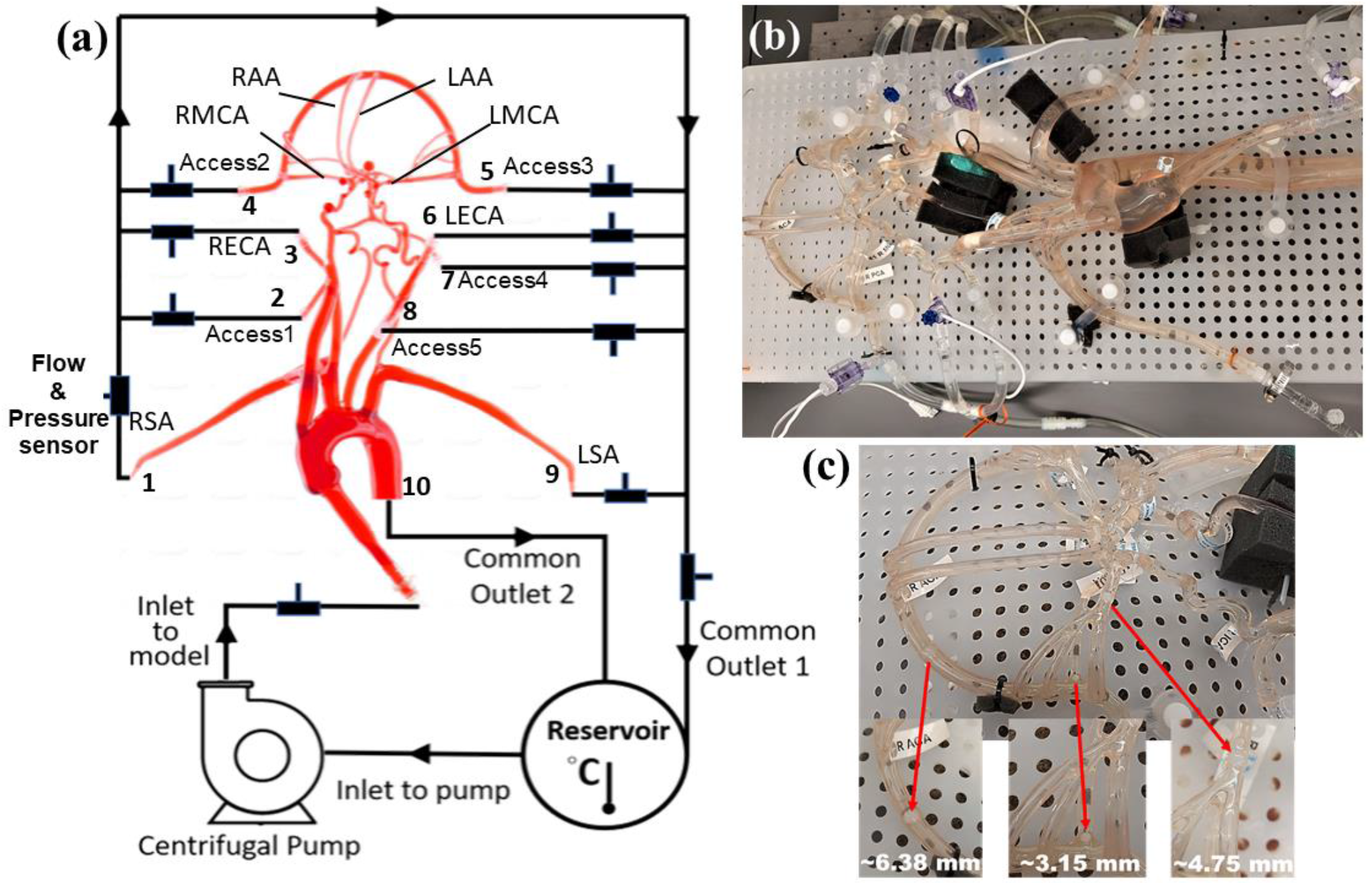
Schematic and photographs of the in vitro circulatory flow loop illustrating the (a) experimental setup, (b) the flow loop setup with the instrumentation that was used to measure flow rate and pressure at various cerebrovascular outlets, and (c) location of nylon spherical clots used to mimic a stroke condition. Here, LSA and RSA are the left and right subclavian arteries, LECA and RECA are the left and right external carotid arteries, LMCA and RMCA are the left and right middle cerebral arteries, and LAA and RAA are the left and right anterior arteries, respectively.

**Figure 2.**
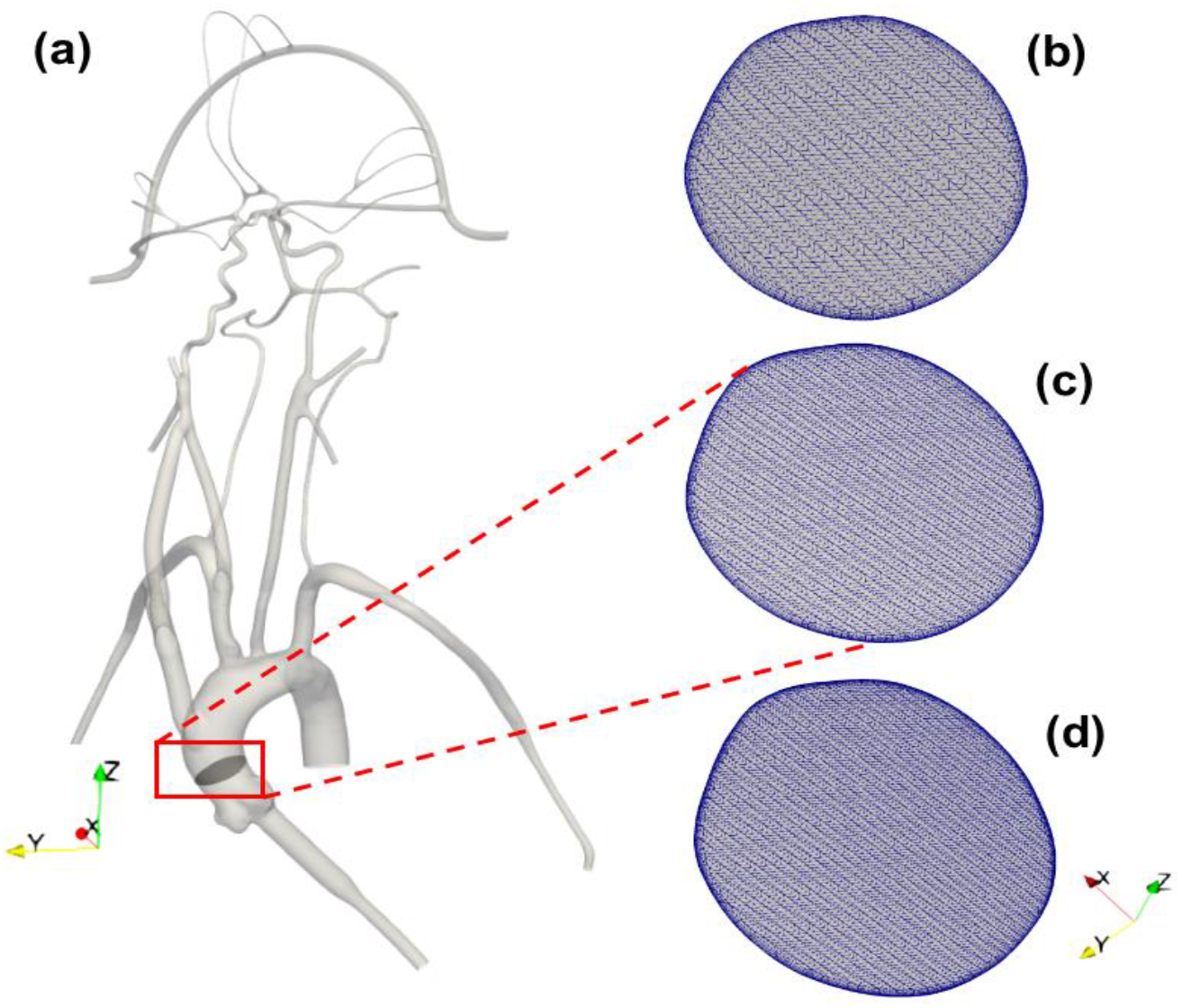
(a) 3D reconstructed computational model of the corresponding *in vitro* cerebrovascular model with a cut plane view in the aortic arch to show the mesh for the (b) 3 M, (c) 6.5 M and (d) 9 M cell meshes.

### 2.2. *In vitro* experiments

A closed-loop circulatory flow system was designed that included a centrifugal pump (Cole-Parmer, IL, USA), reservoir, flow meter and ultrasonic flow probes, and pressure transducers as shown in Fig. 1a and 1b. The flow loop was connected to the cerebrovascular model, which was placed in the supine orientation. The inlet and outlet flow rates and pressures were monitored using ultrasonic flow probes (Transonic Systems, Inc., Millis, MA) and pressure transducers (Merit Medical, South Jordan, UT), respectively. The pressure transducers were connected to an analog data acquisition module (DAQ, National Instruments, Texas, USA) and recorded using LabVIEW software (National Instruments, Texas, USA).

Similar to Riley et al. [26], the working fluid consisted of a mixture of water (60% by weight) and glycerol (40% by weight) to obtain a density and dynamic viscosity that is representative of blood (1.09 ± 0.03 g/mL and 3.98 ± 0.14 cP, respectively) at an operating temperature of 22.2 °C. Experiments were performed using a steady inlet flow rate of 5.17 ± 0.078 L/min, corresponding to a Reynolds number (*Re*) in the inlet tube of approximately 3890. This inlet flow rate was chosen to correspond to a representative mean physiological cardiac output of an adult. An extended tube of 900 mm in length was attached to the model inlet such that the flow entering the model inlet was fully developed. To study the effect of a stroke condition on the mean arterial pressure and flow rate in the cerebral arteries, nylon spherical clots of three different sizes (3.15 mm, 4.75 mm, and 6.38 mm in diameter) were manually inserted into the right MCA (RMCA) to completely block the vessel and the corresponding flow through outlet 4, as depicted in Fig. 1c.

Three separate experiments were performed for both conditions (normal and stroke) to measure the flow rate and pressure at the inlet and various outlets in order to provide boundary conditions for the CFD simulations and for validation. Because the fluid heats up as it is continuously pumped through the flow loop, we waited for 30 minutes to allow for the fluid temperature to reach a steady state (22.2 °C) before measuring the flow rate and pressure. Importantly, the viscosity of the fluid was tuned to account for the effect of this temperature rise, so as to obtain the desired value of 3.98 cP at the steady-state operating temperature. The regional distribution of the flow rate was tuned to match available literature data [30-31] by adjusting clamps downstream of the pressure and flow rate measurement sites. This yielded a regional flow distribution such that 73.2% of the flow passed through the descending aorta and 26.8% of the flow was distributed to the remaining arteries stemming from the aortic arch. The flow rates to individual arteries were also tuned to match that reported in the literature [30-31], which are summarized in Table 1.

**Table 1:**
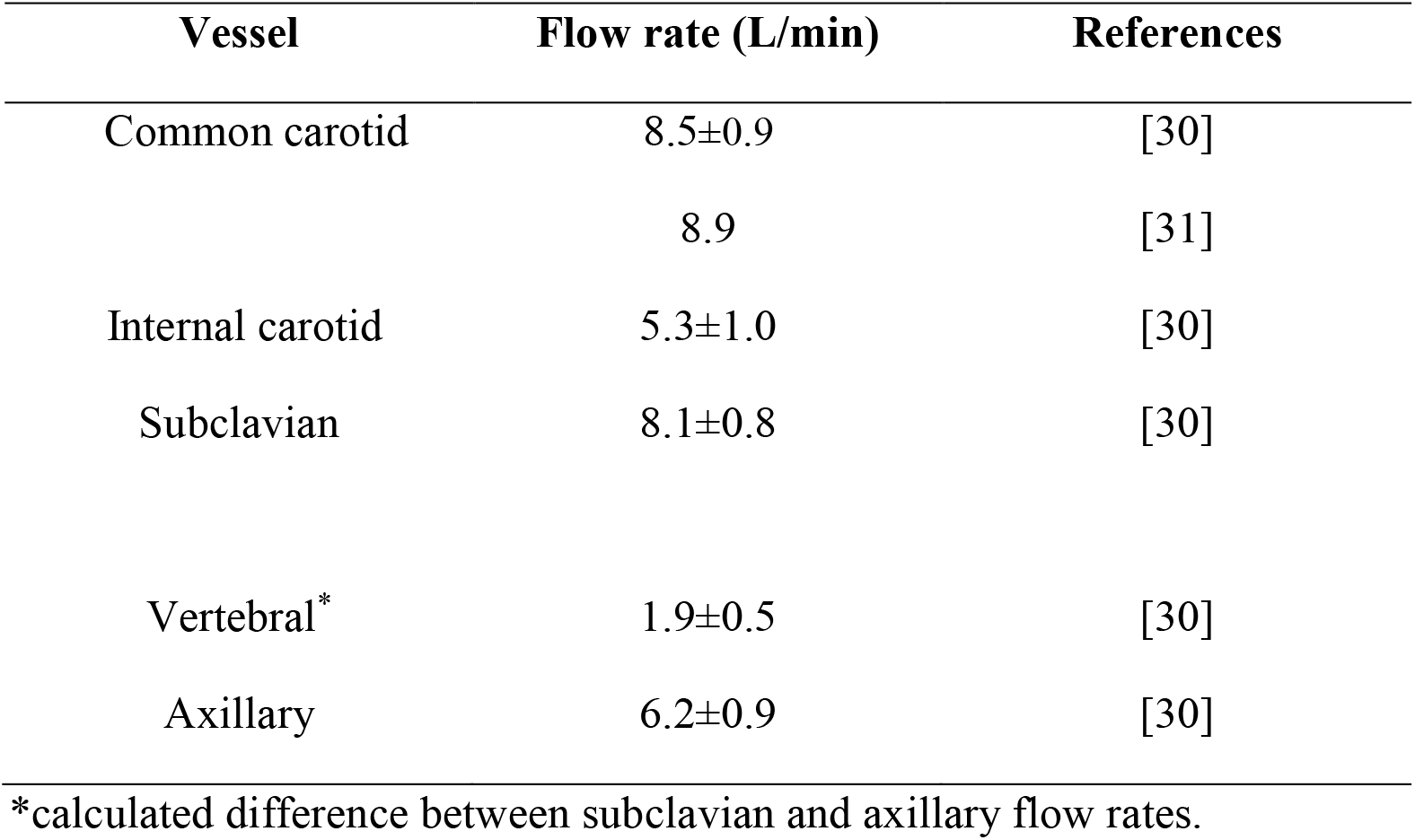
Distribution of cerebral volumetric flow rates from the literature. The flow rate values are reported as mean ± standard deviation (SD).

| Vessel | Flow rate (L/min) | References |
| --- | --- | --- |
| Common carotid | 8.5±0.9 | [30] |
|  | 8.9 | [31] |
| Internal carotid | 5.3±1.0 | [30] |
| Subclavian | 8.1±0.8 | [30] |
| Vertebral* | 1.9±0.5 | [30] |
| Axillary | 6.2±0.9 | [30] |

\*calculated difference between subclavian and axillary flow rates.

### 2.3. CFD mesh

The computational geometry was discretized in CF-Mesh+ (version 3.5.1; Creative Fields, Croatia) using an unstructured hexahedral mesh with 5 wall-normal inflation layers at the walls to resolve large velocity gradients. Three different resolutions of meshes were generated yielding coarse, medium, and fine meshes having 3 million, 6.5 million, and 9 million computational cells, respectively (Figure 2). These generated meshes were used to perform a mesh refinement study to evaluate the sensitivity of the results to the mesh resolution. Using the *topoSet* utility in OpenFOAM, the mesh for the stroke condition was generated by removing the cells covering the section of the arteries that were blocked in the experiments by clots. After eliminating the cells at the clot insertion sites, the newly created surfaces were converted to walls.

### 2.4. Governing equations & boundary conditions

Using the 3D reconstructed model, CFD simulations of the flow were performed by solving the Reynolds-averaged Navier-Stokes (RANS) equations using the open-source computational continuum mechanics library, OpenFOAM (version 2106). The gravitational force is not considered in the present simulations. The effects of turbulence were modeled using the two-equation eddy-viscosity k-ω shear stress transport (SST) turbulence model [32-33]. The k-ω SST model was chosen because of its reported good behavior in adverse pressure gradients and separated flows [34], and because it has been shown to be capable of handling multi-regime (laminar, transitional, turbulent) flows [35].

The CFD boundary conditions were specified to match the corresponding *in vitro* experiments. A constant, uniform inlet velocity was applied to the extended inlet to obtain fully developed flow entering the inlet to the ascending aorta at a flow rate of 5.17 L/min. A no-slip velocity boundary condition was applied on the walls, which were assumed to be rigid. The *pressureInletOutletVelocity* condition in OpenFOAM was applied for the velocity on all outlet boundaries, which uses a zero-gradient Neumann condition on boundary faces with outflow and an extrapolated Dirichlet condition on faces with reversed inflow. A zero-gradient pressure condition was applied at the inlet and fixed static pressure boundary conditions were prescribed on all outlets. The values of outlet pressure were specified by iteratively performing simulations to match the measured flow rate through each outlet in the experiments to within ±10%.

### 2.5. Numerical methods

Simulations were performed in OpenFOAM using second-order accurate discretization schemes, including second-order upwind for the advective terms. Preliminary steady simulations were performed by solving the steady RANS equations of motion using the semi-implicit method for pressure-linked equations (SIMPLE) solver, *simpleFoam*. The steady simulations did not converge owing to regions of significant unsteady flow in the ascending aorta and the aortic arch. Fully transient flow simulations were performed using the hybrid pressure-implicit with splitting of operators (PISO)-SIMPLE solver, *pimpleFoam*. An initial unsteady simulation was performed for 4 seconds of physical time, which was determined to be the time that was needed for the flow to overcome the initial startup transient conditions and to obtain a stationary, quasi-steady flow state. Simulations were then run for another 4 seconds at these quasi-steady conditions while time-averaging the flow solution. The simulations were performed on 40 compute cores of a high-performance computing (HPC) system at the Institute for Computational and Data Sciences at the Pennsylvania State University. Post-processing and visualization of the simulation results were performed in ParaView (Kitware, Inc., Clifton Park, NY).

## 3. Results

### 3.1. *In vitro* experiments

The values of volumetric flow rate (L/min) and pressure (mmHg) measured at various arterial outlets are shown in Figure 3a. As mentioned in Section 2.2, the arteries stemming from the aortic arch, which includes the axillary, external, middle, anterior, and other arteries, receive 26.8% of the total flow rate entering the model and 73.2% of the fluid flows through the descending aorta. Out of the total flow rate, 10.23% travels through both axillary arteries, 6.02% through both external carotid arteries (ECA), and 5.62% through the middle and anterior cerebral arteries. This measured flow distribution matches physiological values from the available literature [30-31]. However, it is important to note that these distributions are based on the steady flow measurements, whereas the values reported in the literature are based on total volumetric flow rate values over a pulsatile cardiac cycle.

**Figure 3.**
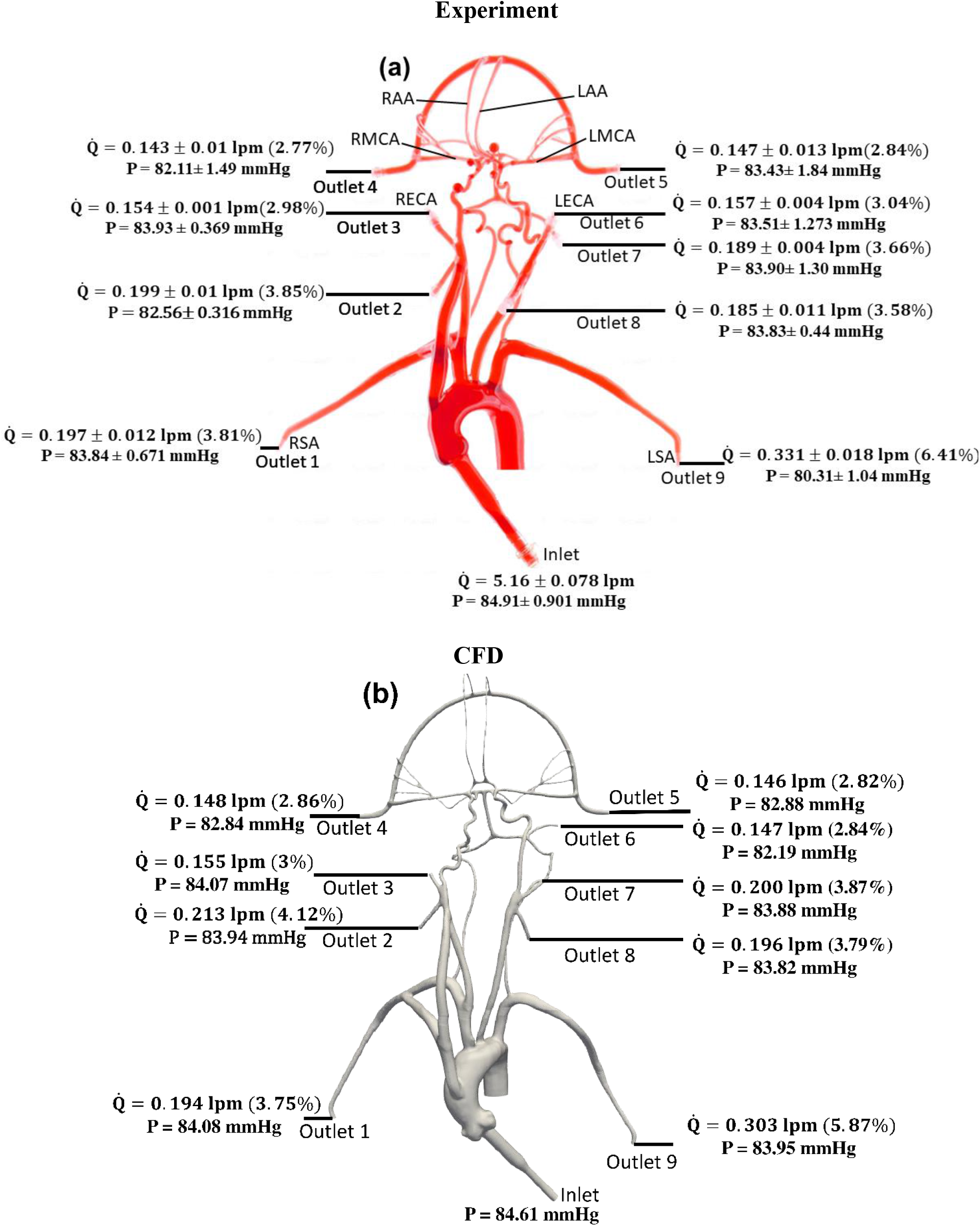
Mean volumetric flow rate and pressure values along with standard deviation at the inlet and all arterial outlets from (a) *in vitro* experiments and (b) CFD simulation for the normal condition. The percentage values shown in parentheses indicate the fraction of total inlet flow through each arterial outlet.

Changes in mean arterial pressure and volumetric flow rate were also measured for the stroke condition when there is no flow from outlet 4. The measurements showed that the mean pressure at all outlets decreased for the stroke condition. As illustrated in Figs. 3 and 4, the flow rate through the right arterial outlets as well as outlets 5 and 6 increased to compensate for the flow that was obstructed due to the emulated stroke.

**Figure 4.**
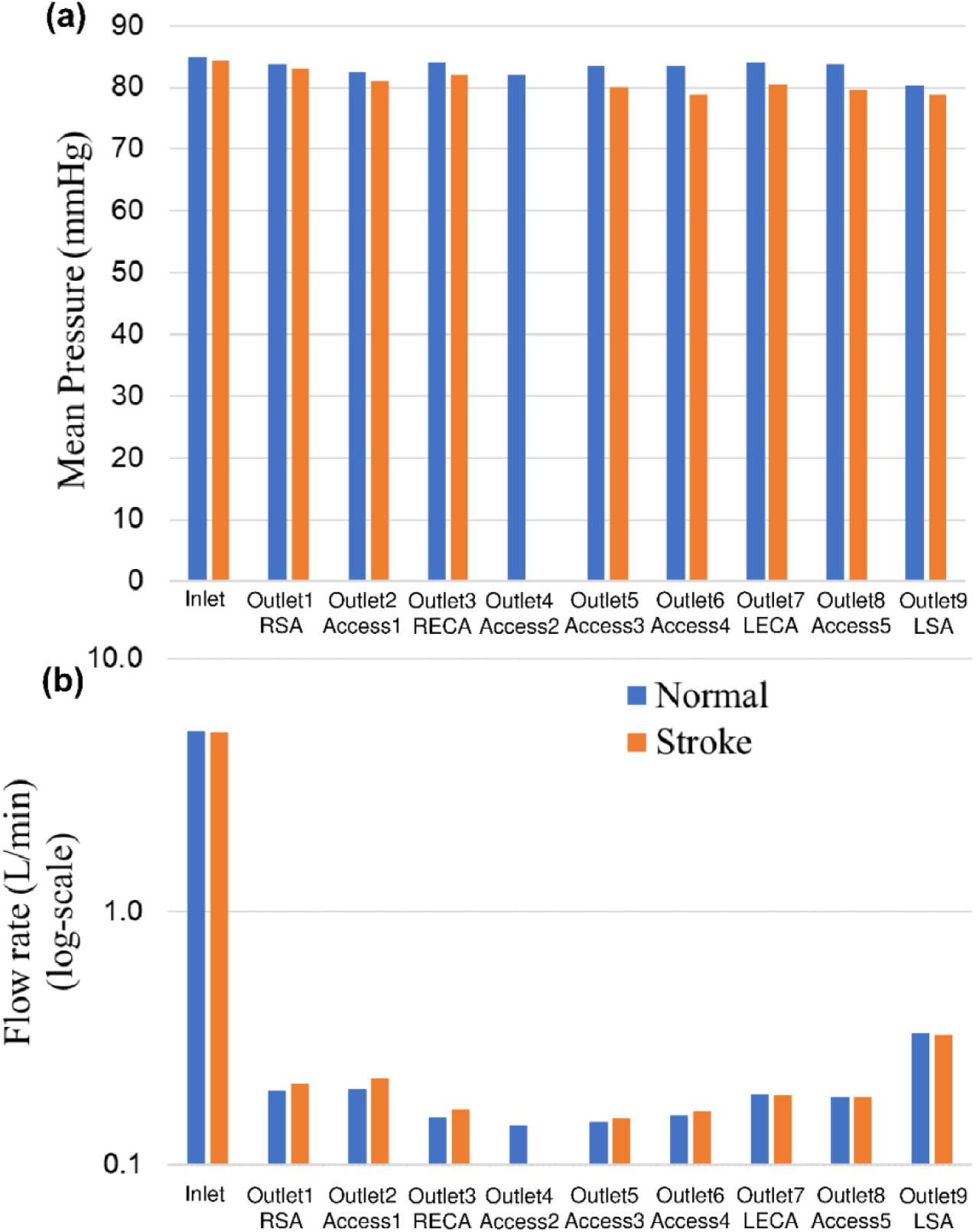
Comparison of (a) mean pressure and (b) flow rate (log scale) at the inlet and various aterial outlets for the normal and stroke conditions from in vitro experiments. The values of pressure and flow rate at outlet 4 for the stroke condition are zero because there is no flow due to the blockage.

### 3.2. CFD flow simulation

To verify that the k-ω SST turbulence model is behaving as expected and producing turbulent flow in regions of expected turbulence and laminar flow in the downstream cerebral arteries, we calculated the regional distribution of the turbulent viscosity ratio (ν_*t*_/ν) that is defined as the ratio of turbulent viscosity (ν_*t*_) to the molecular viscosity (ν). In regions where the k-ω SST model predicts turbulence, ν_*t*_/ν is much larger than unity and in regions of laminar flow ν_*t*_/ν is much less than 1. As illustrated in Fig. 5, the k-omega SST model predicts turbulent flow in the inlet and in the aorta where the flow is expected to be turbulent. In the downstream cerebral arteries, we observe that the predicted flow laminarizes, as evidenced by the negligible turbulent viscosity ratio, indicating that the turbulence model is inactive in this region, yielding laminar flow as expected.

**Figure 5.**
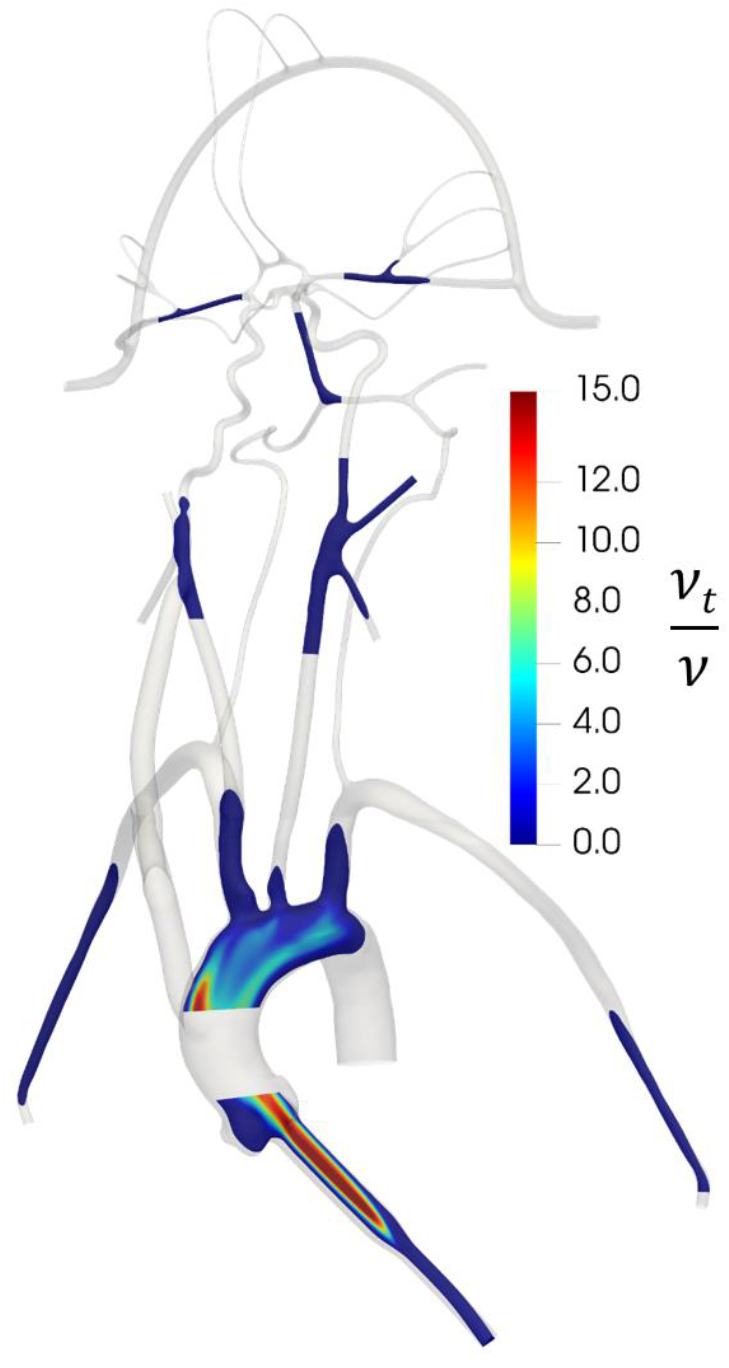
Contours of the distribution of the ratio of turbulent viscosity (ν_*t*_) to molecular viscosity (ν) in the CFD model, illustrating regions of predicted turbulent flow where ν_*t*_/ν is large and regions of laminar flow where ν_*t*_/ν is small.

To evaluate the sensitivity of the CFD results to the mesh resolution, we performed a mesh refinement study to compare the results from the coarse, medium, and fine meshes that contain 3 million, 6.5 million, and 9 million computational cells, respectively. For the same inlet flow rate and with the same outlet pressures applied to each model, we compared the predicted volumetric flow rate through each outlet for the normal physiological condition. The percent differences in the volumetric flow rate at outlet 1, outlet 3, outlet 5, outlet 6, and outlet 9 were calculated to be approximately 1%, 2%, 0.3%, 0.4%, and 0.3%, respectively, between the medium and fine meshes. Between the coarse and medium meshes the percent differences for these same outlets were approximately 3%, 9%, 2%, 3%, and 1%, respectively. Thus, the results between the medium and fine meshes are in close agreement and, therefore, we chose the medium mesh for the final simulations of both the normal and stroke conditions.

Figure 6 illustrates pressure contours and velocity profiles in the cerebrovascular model from CFD for the normal condition. As shown, higher pressures are observed in the aorta compared to the cerebral arteries due to the fact that most of the pressure drop occurs in the smaller cerebral arteries. As the aortic root is attached with a smaller diameter inlet tube, there is some unsteadiness and recirculating flow in the downstream ascending aorta. Generally, the highest velocities are observed in the inlet tube. Flow speeds in the aortic arch are also much higher compared to the more distal downstream arteries. Additionally, it is interesting that the simulation revealed that the flow in the left common carotid artery has a higher velocity than that in the right common carotid.

**Figure 6.**
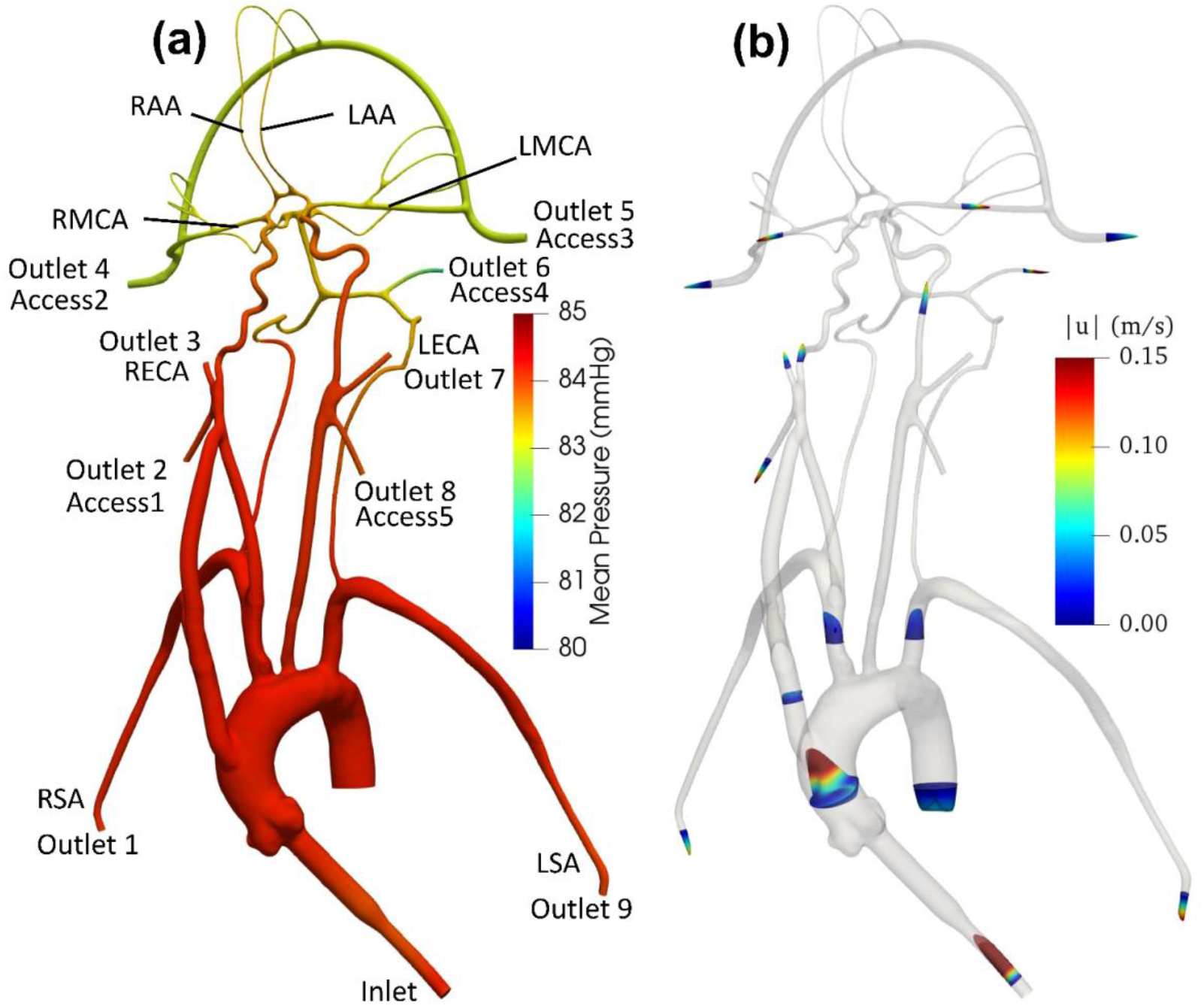
(a) Mean pressure contours and (b) profiles of mean velocity at various sections in the anatomical cerebrovascular model from CFD for the normal condition.

Quantitative values of volumetric flow rate and pressure at all of the outlets from the CFD simulation at the normal condition are summarized in Figure 3b. The flow distribution in the axillary, external, and combined middle and anterior arteries are equal to 9.62%, 6.87%, and 5.68%, respectively. Because outlet boundary conditions were assigned to match the experimental flow rate measurement through each artery outlet to within ±10%, we validate the simulations by comparing the resultant outlet pressures from CFD with the experimental pressure measurements. As summarized in Tables 2 and 3, all of the arterial outlet pressures from CFD are close to the experimental values to within ±7%, with many of the CFD outlet pressures within the uncertainty of the experimental measurement.

**Table 2:**
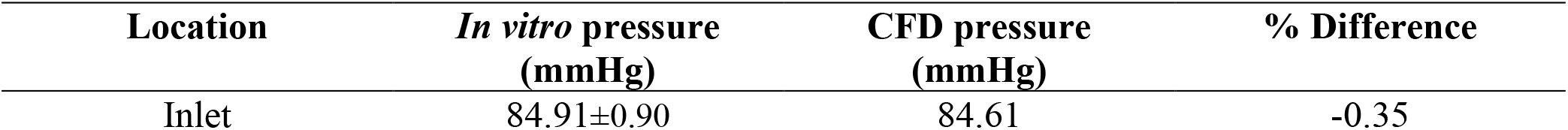
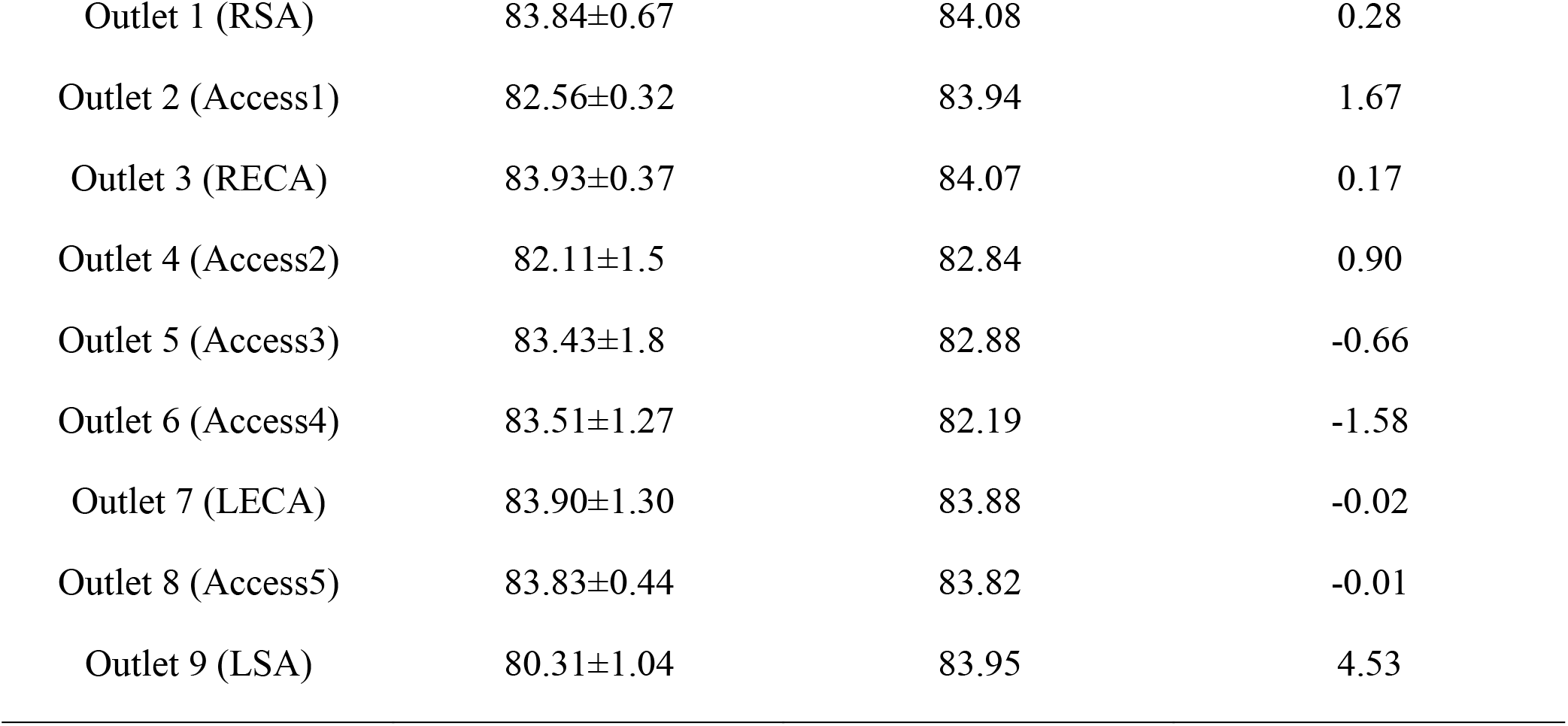
Mean pressure (mmHg) obtained from *in vitro* experiments and CFD at various inlet and outlets for the normal condition. The experimental values are reported as mean ± standard deviation (SD) from three experiments.

| Location | *In vitro* pressure (mmHg) | CFD pressure (mmHg) | % Difference |
| --- | --- | --- | --- |
| Inlet | 84.91±0.90 | 84.61 | -0.35 |
| Outlet 1 (RSA) | 83.84±0.67 | 84.08 | 0.28 |
| Outlet 2 (Access1) | 82.56±0.32 | 83.94 | 1.67 |
| Outlet 3 (RECA) | 83.93±0.37 | 84.07 | 0.17 |
| Outlet 4 (Access2) | 82.11±1.5 | 82.84 | 0.90 |
| Outlet 5 (Access3) | 83.43±1.8 | 82.88 | -0.66 |
| Outlet 6 (Access4) | 83.51±1.27 | 82.19 | -1.58 |
| Outlet 7 (LECA) | 83.90±1.30 | 83.88 | -0.02 |
| Outlet 8 (Access5) | 83.83±0.44 | 83.82 | -0.01 |
| Outlet 9 (LSA) | 80.31±1.04 | 83.95 | 4.53 |

**Table 3:** Mean pressure (mmHg) obtained from *in vitro* experiments and CFD at the inlet and outlets for the stroke condition. The experimental values are reported as mean ± standard deviation (SD) from three experiments.

| Location | *In vitro* pressure<br>(mmHg) | CFD pressure<br>(mmHg) | % Difference |
| --- | --- | --- | --- |
| Inlet | 84.25±0.79 | 84.64 | 0.46 |
| Outlet 1 (RSA) | 83.04±0.85 | 84.09 | 1.26 |
| Outlet 2 (Access1) | 81.01±0.72 | 83.84 | 3.49 |
| Outlet 3 (RECA) | 82.02±0.67 | 84.04 | 2.46 |
| Outlet 4 (Access2) | - | - | - |
| Outlet 5 (Access3) | 80.04±1.2 | 83.13 | 3.86 |
| Outlet 6 (Access4) | 78.83±1.7 | 82.19 | 4.26 |
| Outlet 7 (LECA) | 80.40±1.19 | 83.96 | 4.42 |
| Outlet 8 (Access5) | 79.55±0.96 | 83.89 | 5.45 |
| Outlet 9 (LSA) | 78.74±1.56 | 83.94 | 6.60 |

## 4. Discussion

The objective of this study is to establish a realistic *in vitro* experimental model and a corresponding computational model of the cerebral vasculature that can be used to investigate flow and embolus migration and lodging in the brain. Measurements of pressure and flow rate were acquired and corresponding CFD simulations were performed and compared with the experiments to validate the flow predictions. Overall, the CFD predictions were in relatively close agreement with the experiments, to within ±7% of the mean experimental pressure measurements with many of the CFD predictions within the uncertainty of the experimental measurement.

The *in vitro* model developed in this study represents a realistic anatomical model of the cerebrovasculature and the upstream arteries that produce a realistic flow distribution. The measurements of the flow distribution are in close agreement with the range of clinical values reported in the literature [30-31]. Compared with other *in vitro* cerebrovascular models, the present model consists of the major cerebral vasculature and the aorta. In contrast, previous work (e.g., [36-38]) used models that did not incorporate all of the vasculature that is considered in the present study.

The present CFD simulations closely correspond with the *in vitro* experimental measurements of pressure in the various arterial outlets of the model. Interestingly, as found in previous studies [39-40], we observed that the regional flow distribution in the cerebrovascular model was highly sensitive to the prescribed outlet pressures. In an exploratory sensitivity study, it was observed that small variations in the prescribed outlet pressures within the range of the uncertainty of the experimental pressure measurements yielded extremely large variations in the regional flow distribution in the model. Also, the flow pattern in the aortic arch in the current investigation generally agrees with that reported in the computational study of Numata et al. [41]. The transient flow field found in the aortic arch in the present work has also been reported in other studies [42-45].

Finally, there are several limitations of the present study that should be addressed in future work. First, only flow is measured and simulated in the model; we do not consider embolus migration and lodging, which is a topic of ongoing research. Additionally, steady flow is considered in this study, whereas the physiological flow in the aorta and cerebrovascular is pulsatile in nature. The objective of the current research, however, is to provide a tiered validation data set for systematically validating computational simulations, the first step of which is to validate by comparing with steady flow. Also, the working fluid used in this study is Newtonian. The non-Newtonian rheology of blood is planned to be included in future work. Finally, the CFD simulations in this study assume that the walls of the vascular model are rigid, which is a reasonable assumption for steady flow through the model. Future work investigating pulsatile flow, however, could also consider the influence of vessel wall motion due to fluid-structure interaction and its influence on the flow and migration of emboli.

## Nomenclature

CCA: Common carotid artery
ICA: Internal carotid artery
ECA: External carotid artery
LECA: Left external carotid artery
RECA: Right external carotid artery
MCA: Middle cerebral artery
LMCA: Left middle cerebral artery
RMCA: Right middle cerebral artery
ACA: Anterior cerebral artery
LAA: Left anterior artery
RAA: Right anterior artery
LSA: Left subclavian artery
RSA: Right subclavian artery

## Acknowledgements

We thank Kenneth Aycock for reviewing the manuscript. This work was partially supported by the NIH (NHLBI) through Grant HL146921. NIH did not have any involvement in the study design, in the collection, analysis and interpretation of data; in the writing of the manuscript; and in the decision to submit the manuscript for publication. The findings and conclusions in this article have not been formally disseminated by the U.S. FDA and should not be construed to represent any agency determination or policy. The mention of commercial products, their sources, or their use in connection with material reported herein is not to be construed as either an actual or implied endorsement of such products by the Department of Health and Human Services.

